# Correlation of biomechanics and cancer cell phenotype by combined Brillouin and Raman spectroscopy of U87-MG glioblastoma cells

**DOI:** 10.1101/2022.03.09.483576

**Authors:** Jan Rix, Ortrud Uckermann, Katrin Kirsche, Gabriele Schackert, Edmund Koch, Matthias Kirsch, Roberta Galli

**Affiliations:** Clinical Sensoring and Monitoring, Department of Anesthesiology and Intensive Care Medicine, Faculty of Medicine Carl Gustav Carus, TU Dresden, Fetscherstrasse 74, D-01307 Dresden, Germany; Neurosurgery, Faculty of Medicine Carl Gustav Carus, TU Dresden, Fetscherstrasse 74, D-01307 Dresden, Germany; Division of Medical Biology, Department of Psychiatry, Faculty of Medicine and University Hospital Carl Gustav Carus, TU Dresden, Fetscherstrasse 74, D-01307 Dresden, Germany; Klinik für Neurochirurgie, Asklepios Kliniken Schildautal, Karl-Herold-Strasse 1, D-38723 Seesen, Germany; National Center for Tumor Diseases (NCT), Partner Site Dresden, Fetscherstrasse 74, D-01307 Dresden, Germany; Department of Medical Physics and Biomedical Engineering, Faculty of Medicine Carl Gustav Carus, TU Dresden, Fetscherstrasse 74, D-01307 Dresden, Germany

**Keywords:** Brillouin, Raman, spheroids, glioblastoma

## Abstract

The elucidation of biomechanics furthers understanding of brain tumor biology. Brillouin spectroscopy is a new optical method that addresses viscoelastic properties down to subcellular resolution in contact-free manner. Moreover, it can be combined with Raman spectroscopy to obtain co-localized biochemical information. Here, we applied co-registered Brillouin and Raman spectroscopy to U87-MG human glioblastoma cells in vitro. Using 2D and 3D cultures, we related biomechanical properties with local biochemical composition at subcellular level, as well as cell phenotype. Brillouin and Raman mapping of adherent cells showed that the nucleus and nucleoli are stiffer than the perinuclear region and the cytoplasm. The biomechanics of cell cytoplasm is affected by culturing conditions, i.e. cells grown as spheroids being stiffer than adherent cells. Inside the spheroids, the presence of lipid droplets as assessed by Raman spectroscopy reveals higher Brillouin shifts which is not related to local stiffness increase, but due to a higher refractive index combined with a lower mass density. This highlights the importance of locally defined biochemical reference data for a correct interpretation of the Brillouin shift of cells and tissue in future studies investigating the biomechanics of brain tumor models by Brillouin spectroscopy.

## 1. Introduction

The importance of biomechanics for tumor biology is increasingly acknowledged (1). Tumors generally exhibit biochemical and biomechanical properties that differ from those of normal tissue. In addition, the metastatic potential of tumor cells is linked to the cell’s mechanical properties (2). Softer cell nuclei are related to higher metastatic spread (3). Moreover, recent research suggests that mechanical stress and increased activation of mechanosignaling promote malignant transformation and metastatic processes (4). It also affects tissue perfusion, as well as angiogenesis (1). Comprehensive analysis of biomechanics provides important insights into disease-induced changes in stiffness (5). Therefore, strategies that consider tumor mechanics might lead to effective therapeutic approaches of treatment-resistant or metastatic cancer (6).

In neurooncology, the study of cell biomechanics is still in its infancy, with some research attributing tremendous importance to it. Atomic force microscopy (AFM) allowed discriminating WHO grade II, III and IV astrocytomas and thus between different degrees of malignancy (7). In experimental gliomas and brain metastases, magnetic resonance elastography demonstrated decreased viscosity and elasticity compared to brain parenchyma. In this regard, brain metastases with an infiltrative growth pattern were softer than solid glioma (8). Furthermore, a correlation between the strength of the extracellular matrix and the aggressiveness of brain tumors was established based on a change in mechanosignaling (9) and durotactic stimuli were identified as a major factor for glioma cell migration (10). Structure, motility and proliferation of glioma cells are influenced by biomechanical properties of the tissue (11). However, there appears to be high interpatient variability with respect to responses to biomechanical stimuli, as shown in experiments with primary glioblastoma cell lines (12). Systematic studies addressing cellular and subcellular biomechanical properties of brain tumor cells are lacking so far.

The mechanics of cells can be determined by various methods, however AFM is currently most commonly used in tumor research (13). Several studies performed with AFM show that cancer cells can be distinguished from normal tissue, as well as original and metastatic cancer cells, by analyzing their mechanical properties. Metastatic carcinoma cells were identified by mechanical studies using AFM (14,15). Furthermore, several authors reported tumor cells being generally softer than normal cells (16–18). However, all studies with AFM have an inherent problem arising from the contact between the measuring instrument and the sample: Since AFM probes the surface of the sample, just information about the biomechanics of the cell as a whole is obtained.

As an alternative optical contact-free technique, Brillouin spectroscopy exploits the inelastic scattering of photons of a laser beam upon interaction with GHz-frequency acoustic phonons in the sample. This technique is used to determine the elastic properties of materials by probing the frequency of the Brillouin shift. Brillouin spectroscopy avoids any contact with the sample while providing biomechanical information at subcellular level (19). The Brillouin shift describes viscoelasticity and is thus not equivalent to an analysis of rigidity (Young’s modulus) that is obtained by AFM. Furthermore, the Brillouin shift depends also on the local index of refraction *n* and mass density *ρ*, which shall be known to retrieve the longitudinal elastic modulus. Nevertheless, it has been shown that Brillouin spectroscopy can reveal local changes in biomechanics of cells and tissues also without a priori knowledge of *n* and *ρ* (20,21), and that changes in Brillouin shift correlate with changes in Young’s modulus (20,22). Therefore, the Brillouin shift is considered a proxy for the stiffness of biological materials.

Brillouin spectroscopy became available only recently for biomedical applications because of unsolved technical challenges. The Brillouin shift is extremely small (< 0.001 nm), thus separating the Rayleigh scattering from the Brillouin scattering and the spectral analysis of the latter is difficult when measuring turbid materials like biological samples. Only in 2016, Fiore et al. described a relatively simple approach with a spectrograph for Brillouin spectroscopy, encompassing a multi-pass Fabry-Pérot interferometer as ultra-narrow bandpass filter and a highly dispersive optical element obtained by combining two virtually-imaged phased arrays (VIPA) (23). This enabled near-lossless optical isolation of the Brillouin signal that opened the avenue to the study of biological tissue.

Recently, Brillouin microscopy was applied to 3D spheroids which simulate tumors in a more realistic way than single cells by accounting for cell-cell contacts, diffusion gradients, proliferation rates and drug responses (24–26). Studies on colorectal tumor spheroids showed that the mechanical properties were altered heterogeneously across the spheroid after drug treatment (27). Moreover, it was shown that the osmolality of surrounding medium affects the biomechanics of ovarian cancer cell spheroids (22). Furthermore, the effect of micro-environmental stiffness and degradability of hydrogels on breast cancer spheroids was demonstrated (28).

The combination of Brillouin spectroscopy and microscopic setups allows high resolution mapping of biomechanical properties. Thanks to tight laser beam focalization coupled with confocal detection, small spot sizes (~1 μm) are achieved, enabling analysis of distinct cell compartments, e.g., cytoplasm, nucleus, and nucleolus (29,30). It should be mentioned that even smaller spot sizes are not useful as the acoustic phonon wavelength limits further resolution increase (31). Furthermore, Brillouin spectroscopy can be easily combined with Raman spectroscopy for simultaneous chemo-mechanical spectroscopic analysis (32). As Raman spectroscopy allows to draw conclusions about the biochemical composition of a sample (i.e. lipids, proteins, nucleic acids), it may be used as a reference to correlate biochemistry and biomechanics at the very same measurement position. The benefit of a combined measurement has been already demonstrated in some studies including single cells (32–34), human epithelial (35,36) and corneal (37) tissue as well as transgenic mouse hippocampus (38). Besides the general feasibility of combined measurements, it was shown that the biochemical information present in the Raman spectra can be used for interpreting the Brillouin shift (39,40).

The possibility to perform Brillouin spectroscopy with near infrared excitation in order to avoid photodamage and attain much larger penetration depth on bulk samples was demonstrated for in vivo measurement by Schlüßler et al. in 2018, who performed Brillouin spectroscopy of the spinal cord of living zebrafish larvae by using a VIPA-based spectrometer and excitation at 780 nm (41). However, combined systems exploiting near infrared excitation found so far only limited application. Also VIPA-based spectrometers are rarely used in combined systems, where predominantly Fabry-Pérot-based spectrometers were utilized in combination with 532 nm excitation wavelength (32,38,42). However, the sequential spectrum acquisition of Fabry-Pérot-based spectrometers comes along with long acquisition times due to the scanning through the frequencies (43).

Here, we addressed the relationship between Brillouin spectrum and cellular components in brain tumor models in vitro by using a combined Brillouin and Raman confocal microscopic system with near-infrared laser excitation to avoid any photodamage. Aim of this study is the correlation of biomechanics with cancer cell phenotype and biochemical properties of glioma cells cultured in different conditions. Therefore, we addressed adherent and spheroid cell culture preparations of U87-MG glioblastoma. First, we used Raman spectroscopy to identify the cell compartments and used this information to determine the Brillouin shift of each compartment, showing that these have different viscoelastic properties. Then, we addressed the comparison between adherent (2D) and spheroid (3D) cultures of U87-MG glioblastoma cells, showing that the Brillouin shift of spheroid cells is globally higher than of adherent cells. Furthermore, the combination of Raman and Brillouin spectroscopy highlighted biochemical cues underlining the local variation of Brillouin shift observed within the spheroids.

## 2. Materials and Methods

### Cell culture and sample preparation

For adherent cultivation (2D cell preparations), 20,000 - 50,000 human U87-MG cells were seeded on Raman-grade CaF_2_ slides and cultured for two to three days in Dulbecco’s Modified Eagle Medium (DMEM, Thermo Fisher Scientific Inc., Waltham, United States) at 37°C and 5% CO_2_. For Raman-Brillouin mappings, slides were transferred to a dish and completely covered with fresh DMEM. For reference measurements, adherent cells grown on CaF_2_ slides were fixed with 4% formalin for 30 min and washed with aqua dest. twice.

For 3D spheroid cultivation, U87-MG tumor cells were suspended in a cell culture flask and cultured with abundant DMEM for one week. The upright positioning of the culture flask prevented the suspended cells from adhering to the walls, which initially led to the formation of cell clusters in the culture medium, which later on grew up to spheroids. It should be mentioned that no special scaffold was used to enforce any spheroid formation. No hydrogel or scaffold was used but spheroids were cultured free floating in DMEM. Culture flasks were gently shaken to resuspend the spheroids every two to three days. For the measurement, few spheroids were transferred on a CaF_2_ slide and covered with DMEM.

For multiphoton microscopy (MPM), spheroids were collected, embedded in cryotome matrix (OCT, CellPath Ltd., Newtown, United Kingdom) and frozen at −80°C. Cryosections of 10 μm were prepared on glass slides, whereby the fifth section through a spheroid was stored at −20°C until use. CARS imaging of whole spheroids proved homogeneity between internal and external regions, which is in line with previous findings (44).

### Combined Brillouin and Raman spectroscopic system

The layout of the combined Brillouin and Raman system is shown in Figure 1. A tunable diode laser (TApro, TOPTICA Photonics AG, Gräfelfing, Germany) was used as photon source. Stabilization to the rubidium ^85^Rb F_g_ = 3 transition at λ = 780.24 nm ensured high wavelength accuracy by using Doppler-free saturation spectroscopy (CoSy, TOPTICA Photonics AG, Gräfelfing, Germany). Two Bragg gratings (NoiseBlock, ONDAX Inc., Monrovia, United States) filtered out the amplified spontaneous emission (ASE) in order to suppress the background. A doubly passed Fabry-Pérot interferometer (Tunable Fabry-Pérot-Etalon FSR = 15 GHz, LightMachinery Inc., Nepean, Canada) further suppressed the ASE to ensure highest signal-to-noise ratio. In this process, a λ/4-wave plate rotated the light polarization direction so that the light was reflected off the polarizing beam splitter on the return path. One of the ghost beams appearing at the polarizing beam splitter was used to stabilize the Fabry-Pérot interferometer to its maximum transmission by detecting the light power via a photodiode. Another ghost beam was used as excitation light source for a reference beam path.

**Figure 1.**
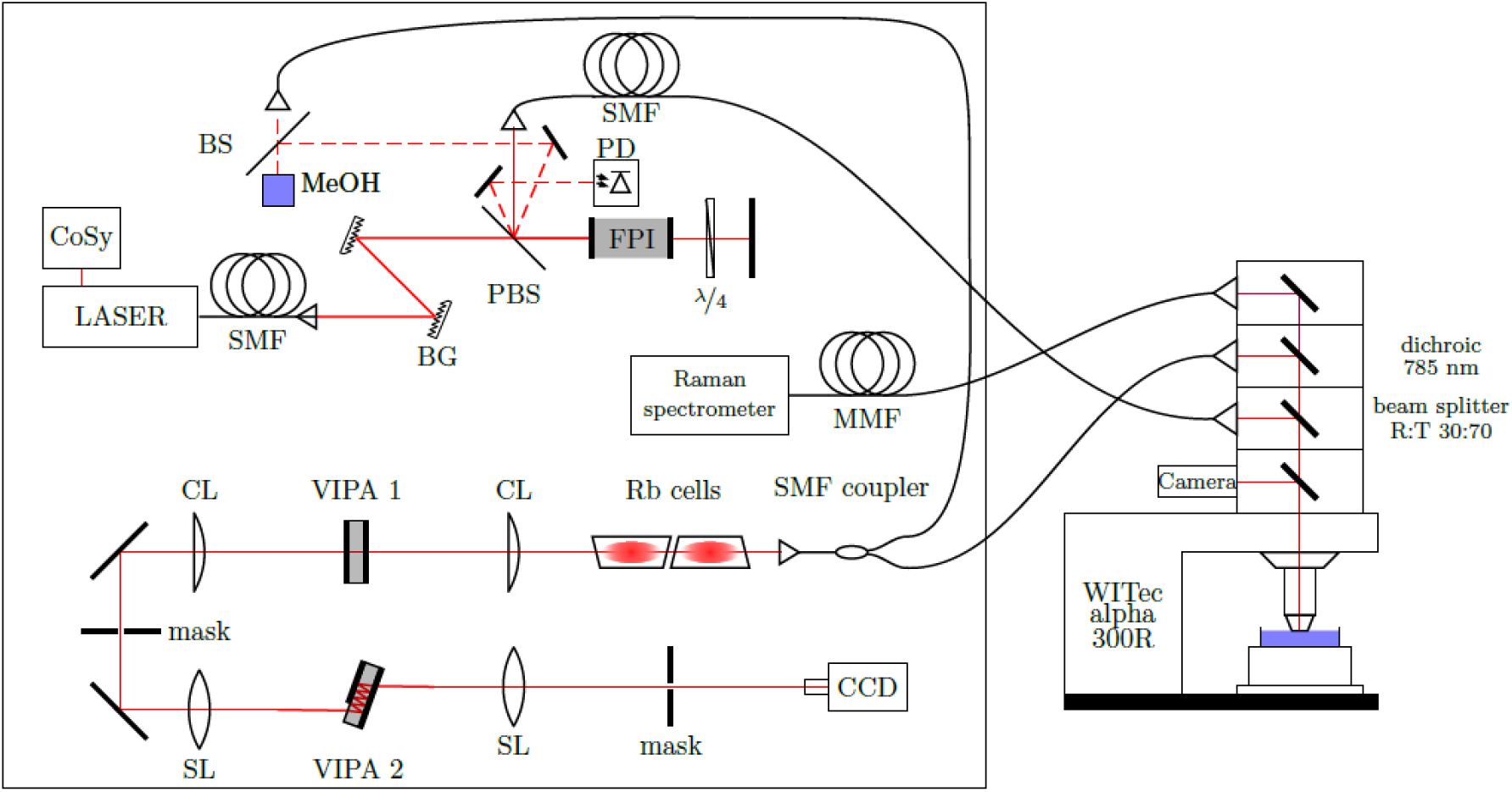
Experimental set-up of the combined Brillouin and Raman system consisting of a 780.24 nm laser, a compact saturation spectroscopy module (CoSy), single/multi-mode fibers (SMF/MMF), Bragg gratings (BG), a polarizing beam splitter (PBS), a Fabry-Pérot interferometer (FPI), a photodiode (PD), neutral beam splitter (BS), cylindrical/spherical lenses (CL/SL) and virtually imaged phased arrays (VIPA).

The excitation light was coupled in a single mode fiber and propagated to an upright reflection microscope (WITec alpha 300R, WITec GmbH, Ulm, Germany). Inside the microscope a 30:70 beam splitter directed the monochromatic light to the sample being placed on a xyz stage. The laser power on the sample was 20 mW. A Zeiss N-Achroplan 40×/0.75NA water-dipping objective was used for adherent cell and spheroid measurements in medium and a Zeiss Epiplan-Neofluar 50×/0.8NA objective for fixed samples.

The backscattered light was separated by wavelength using a dichroic mirror. Light with a wavelength above 785 nm (Raman scattered light) was passed to a commercial Raman spectrometer (UHTS 400, WITec GmbH, Ulm, Germany) whereas light with a wavelength below 785 nm (Brillouin and Rayleigh scattered light) was passed to the Brillouin spectrometer.

In the Brillouin spectrometer, two vapor cells (Rubidium Vapor Cell TG-ABRB-Q, Precision Glassblowing Inc., Englewood, United States) were used to remove the Rayleigh scattered light exploiting absorption on the same rubidium transition used for laser stabilization. A two-stage virtually imaged phased array (VIPA, LightMachinery Inc., Nepean, Canada) set-up (45) was used for the spectral analysis of Brillouin scattering. The orthogonally-arranged VIPAs had a free spectral range of FSR_1_ = 15 GHz and FSR_2_ = 21.6 GHz resulting in a further suppression of the ASE (46). A CCD camera (iDUS 420A-BR-DD, Andor Technology Ltd., Belfast, Northern Ireland) with a magnification objective (InfiniProbe TS-160, Infinity Photo-Optical Company, Centennial, United States) was used to acquire the Brillouin spectra. The detector resolution was 44 MHz/pixel. The optical contrast of the spectrometer (peak-to-background ratio) amounted to 90 dB and the spectral resolution (FWHM of the laser line) was ~400 MHz.

In a reference beam path, the Brillouin signal of methanol was continuously acquired in order to calibrate the spectral axis and thus precisely determine the frequency of the Brillouin band of the measured sample compensating thermally-induced changes.

Acquisition parameters (integration times, accumulations, step sizes) are stated in results section for the different experimental approaches.

### Data analysis

After fitting the Brillouin spectrum with Lorentzian functions using custom-written Matlab software (Matlab, MathWorks Inc., Natick, United States), which is based on the *lsqnonlin* function, the known Brillouin shift of methanol (*ν*_*B*_ = 3.81 GHz (45)) was exploited in combination with the condition that the absolute frequency of the sample’s Stokes and anti-Stokes signal have to be equal. The absolute shift frequency (center), the linewidth (full-width-half-maximum) and the intensity (maximum) of the Brillouin band of the sample were retrieved from the fitting procedure and used to build maps of samples’ biomechanics. Lorentzian functions were used for fitting the Brillouin bands, because they mathematically take the damped-harmonic-oscillator characteristic of an acoustic phonon into account. An exemplary fitted Brillouin spectrum is depicted in the supporting information Figure S1.

For the analysis of single cell Brillouin shift maps, the contribution of surrounding culturing medium was removed in order to focus on the information of the cell itself. Therefore, after fitting the frequency histogram with multiple Gaussian functions (using Matlab’s *lsqnonlin* function), the curve being associated with the medium was subtracted, as it was reported earlier (47,48). Gaussian functions were used, because they take the normal distribution of the measured values into account. It should be mentioned that the elimination of the medium contribution does not affect further analysis of the cells’ mean Brillouin shift, but is rather performed to better visualize the information of interest. Exemplarily, frequency histograms with and without medium contribution are shown in supporting Figure S2.

The Raman spectra were processed according to established protocols for biological samples: Baseline correction followed by intensity normalization were applied (Matlab functions *msbackadj* and *msnorm*). In order to build maps of samples’ chemometric from the Raman data, k-means clustering was used (*kmeans* function of Matlab with squared Euclidean distance metric).

### CARS microscopy and quantification of lipid droplets

The CARS microscope is described in detail elsewhere (49). Briefly, two pulsed Erbium fiber laser (Femto Fiber pro NIR and TNIR, TOPTICA Photonics AG, Gräfelfing, Germany) emitting at a wavelength of 781 nm and 1005 nm were used to resonantly excite the coherent anti-Stokes Raman scattering (CARS) signal of symmetric stretching of the CH_2_ groups, which are mostly contained in lipids. By scanning the lasers over the sample (laser scanning module LSM 7, Carl Zeiss AG, Jena, Germany) the CARS signal is acquired and used to build 2D intensity images (2048 × 2048 pixels, 236 × 236 μm^2^) that enable visualization of lipid droplets within the tissue. Fiji software (43) was used to quantify lipid droplets in CARS images. An empirical determined color threshold of 220 was set for the 8-bit CARS signal to identify areas of high lipid concentration. Afterwards, the *analyze particles* function was used to first determine the number of identified areas with a pixel size greater than 10 and then evaluate the total pixel size of these areas.

## 3. Results

### 3.1. Subcellular compartments of U87-MG glioblastoma cells identified by Raman spectroscopy show different Brillouin shifts

Combined Brillouin and Raman maps were acquired on living U87-MG adherent cells (n = 4, 15 s integration time and averaging of 2 accumulations for each measurement point, 1 μm step size between adjacent points). Figure 2 shows one representative example. The results of the other specimens are shown in supporting information Figure S3.1-S3.3. In Figure 2a the bright field image of the cell is shown and the box indicates the region mapped by spectroscopy.

**Figure 2.**
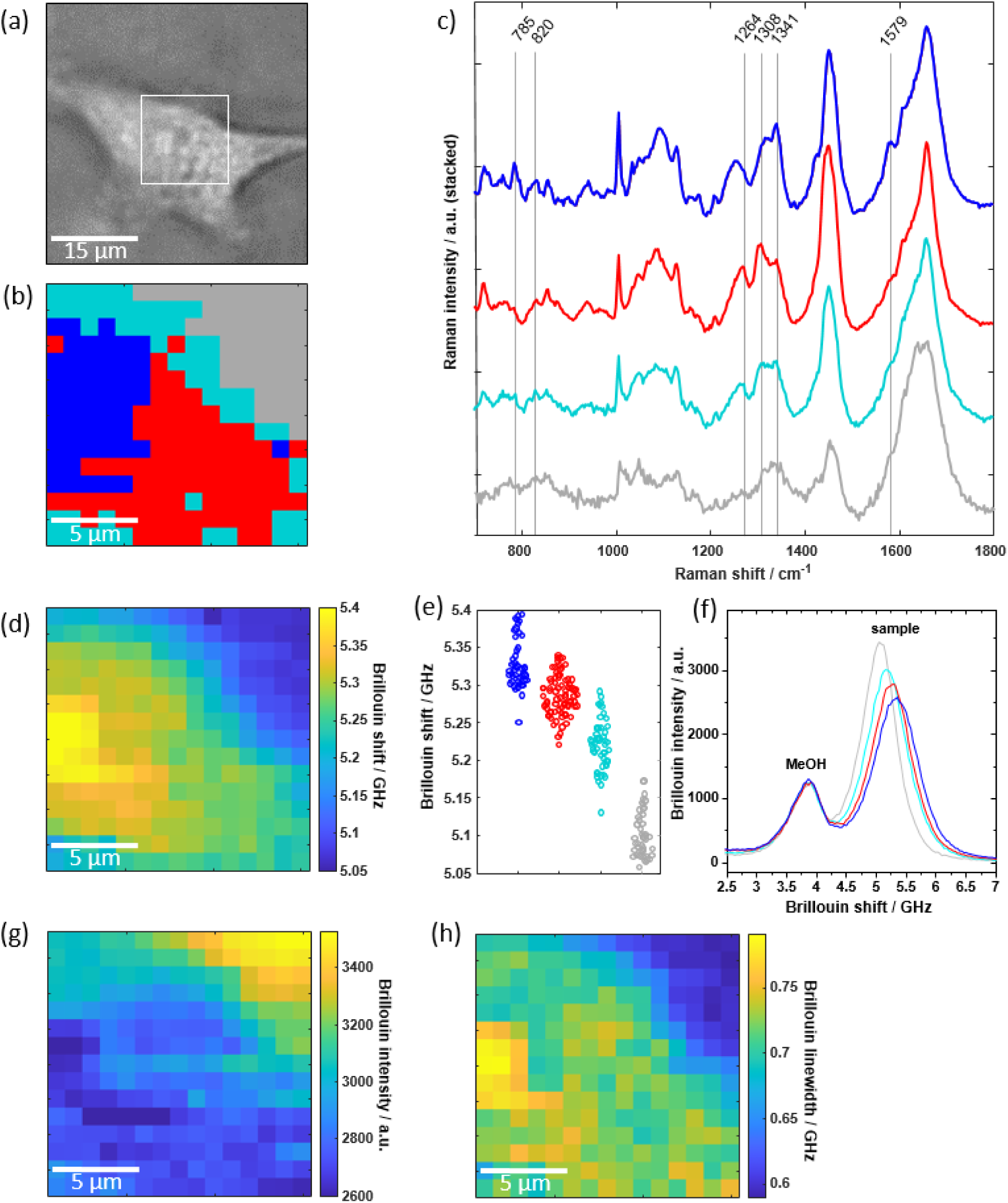
(a) Bright field image of a living U87-MG cell; white box is 15 × 15 μm^2^. (b) Raman cluster map of this cell consisting of four different clusters. (c) Mean Raman spectra of the three clusters associated with the cell being the nucleus (blue), perinuclear region (red) and the cytoplasm (cyan). (d) Simultaneously acquired Brillouin shift and (g) Brillouin intensity map revealing both the cellular structure. (e) Brillouin shift values for each pixel are assigned to the respective cell compartment obtained by the cluster analysis of Raman spectra. (f) Exemplary Brillouin single spectra of each cluster showing the reference methanol band at 3.81 GHz and the sample’s band which shifts to higher values and decreases when going from the gray to the blue cluster. (h) Brillouin linewidth map revealing the high viscosity of the nucleolus.

First, Raman spectroscopy was exploited for identification of mapped regions (Figure 2b). Cluster analysis of the Raman spectra was performed revealing the cell culture medium around the cell (gray cluster) and different cellular compartments. The centroid spectra of the three clusters associated with the cell are depicted in Figure 2c. The blue cluster was attributed to the nucleus with its characteristic DNA bands at 785 cm^−1^, 1341 cm^−1^ and 1579 cm^−1^ (50). The red cluster was associated with a perinuclear region containing characteristic (phospho)lipid bands at 1264 cm^−1^ and 1308 cm^−1^ (50). The cyan cluster was associated with the cytoplasm consisting of different protein bands (e.g. 820 cm^−1^ (50)). The assignments of the clusters to the respective cell compartments are consistent with the measurement positions as indicated in the bright field image. This assignment was also confirmed on fixed U87-MG cells (n = 8), which showed very similar Raman spectra in the same compartments (supporting Figure S4).

The Brillouin shifts plotted as heat map are shown in Figure 2d. A region with higher Brillouin shifts is located in the center of the cell, whereas the cell boundary is characterized by lower Brillouin shifts. By using the results of Raman cluster analysis, the Brillouin shift at the different measurement positions was assigned to the respective cell compartment (Figure 2e). The median Brillouin shift is 5.32 GHz for the nucleus (blue cluster), 5.29 GHz for the perinuclear region (red cluster), 5.22 GHz for the cytoplasm (cyan cluster), and 5.10 GHz for the medium surrounding the cell (gray cluster). Therefore, the highest Brillouin shift was observed in the nucleus, which is consistent with existing literature (47,48). Moreover, it is possible to identify the nucleolus as an intracellular compartment with the highest Brillouin shift.

In Figure 2f, single Brillouin spectra of each cluster are exemplarily shown. The Brillouin band shifts towards higher frequencies by moving from the medium (gray) to the nucleus (blue). Figure 2f also shows that the intensity of Brillouin bands decreases with higher Brillouin shifts. The Brillouin band of methanol used for calibration of the Brillouin shift frequency is visible at 3.81 GHz with constant intensity in all spectra. The Brillouin intensities map is plotted as heat map in Figure 2g and revealed the same morphology as the Brillouin shift. The Brillouin intensity is highest within the culturing medium and decreases inside the nucleus, being lowest in the cell nucleolus. Therefore, both these (inverse correlated (51,52)) parameters enabled mapping of the cell structure.

The width of the Brillouin band was also investigated as proxy for the viscosity (53). A region with highest values is located within the nucleus (Figure 2h). In agreement with (21), this was assigned to the high viscosity of the nucleolus.

### 3.2. Brillouin spectroscopy reveals significantly higher Brillouin shifts for U87-MG spheroids in comparison to U87-MG adherent cells

As acquisition of Raman spectra requires long measurement time, thereby limiting the amount of data that can be acquired on living cells, only the Brillouin spectra were acquired in order to statistically compare the properties of adherent cells and spheroids. Brillouin maps of n = 11 U87-MG adherent cells and n = 9 U87-MG spheroids were acquired using an integration time of 0.2 s (one accumulation for each measurement point, step size of 1 μm for adherent cells and 2 μm for cell spheroids). An example of spheroid mapping is shown in Figure 3. The bright field image of the spheroid is shown in Figure 3a. Corresponding Brillouin maps are depicted in Figure 3b and 3c using the Brillouin shift and the Brillouin intensity as contrast mechanism, respectively. In both maps, which are cross-sections through the spheroid, the spheroid cells can be distinguished from the surrounding culturing medium, which has lower Brillouin frequencies and higher Brillouin intensities. However, not all structures (cf. red arrows) visible in the Brillouin intensity map are observable in the Brillouin shift map and vice versa. This phenomenon might be caused by a sort of shadowing effect, because of absorption and scattering on cells lying above the measurement point, wherefore the Brillouin intensity is reduced. In contrast, the Brillouin shift remains unaffected due to confocal measurement. As the Brillouin intensity is altered in dependence of the absorption of above lying cells, the Brillouin intensity map does not only contain anatomic structures within the plane as reported earlier (41), but rather gives information about the three-dimensional morphological structure.

**Figure 3.**
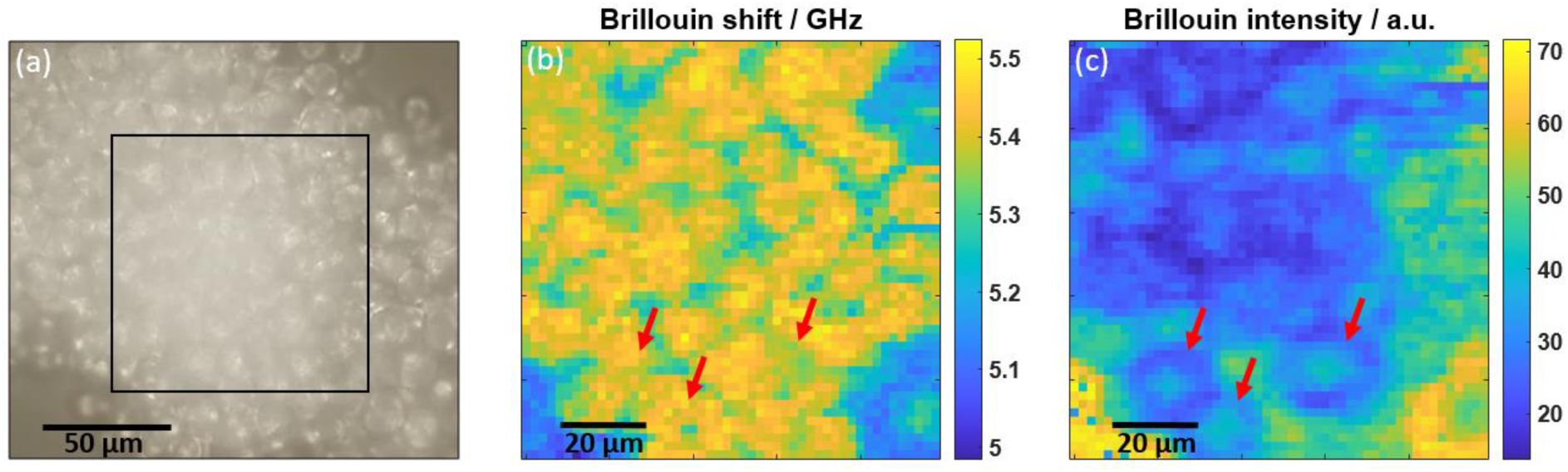
(a) Exemplary bright field image of a U87-MG spheroid, where the black box indicates the measured area. Corresponding Brillouin maps of the same spheroid using (b) the Brillouin shift and (c) the Brillouin intensity as contrast mechanism. Red arrows indicate positions where the structural information between the maps is different to each other.

In order to compare the stiffness of adherent cells and cell spheroids, the Brillouin shifts retrieved from mappings were cumulatively plotted in frequency histograms (Figure 4a and 4b). Here, the data related to culturing medium was eliminated as described in the methods section. Gaussian fitting revealed that the main contributions in the cell and the spheroid maps are different, being the one of spheroid cells located at higher frequency (at about 5.4 GHz vs 5.3 GHz for adherent cells). Additionally, in both histograms a minor contribution is visible at 5.15 GHz, which was associated with the border of cells and is due to a combination of signals from the cell and the medium, based on the results described in the previous section (compare with Figure 2e). Moreover, in the spheroid histogram a minor contribution at 5.34 GHz was associated with the intercellular spaces within the spheroid (compare Brillouin map in Figure 3b). The statistical analysis of the main contributions of the independent measurements showed that the mean Brillouin shift of the cell spheroids is significantly higher than the mean Brillouin shift of the cells (Figure 4c, Mann-Whitney U-test, n = 9 and 11, p < 0.001). The broader distribution in the case of adherent cells is due to a focusing issue, i.e. the Brillouin shift is also dependent on the axial position within a cell. This effect is minor in the case of spheroid maps, because several cells with different axial position are measured, resulting in an averaging and ta smaller distribution.

**Figure 4.**
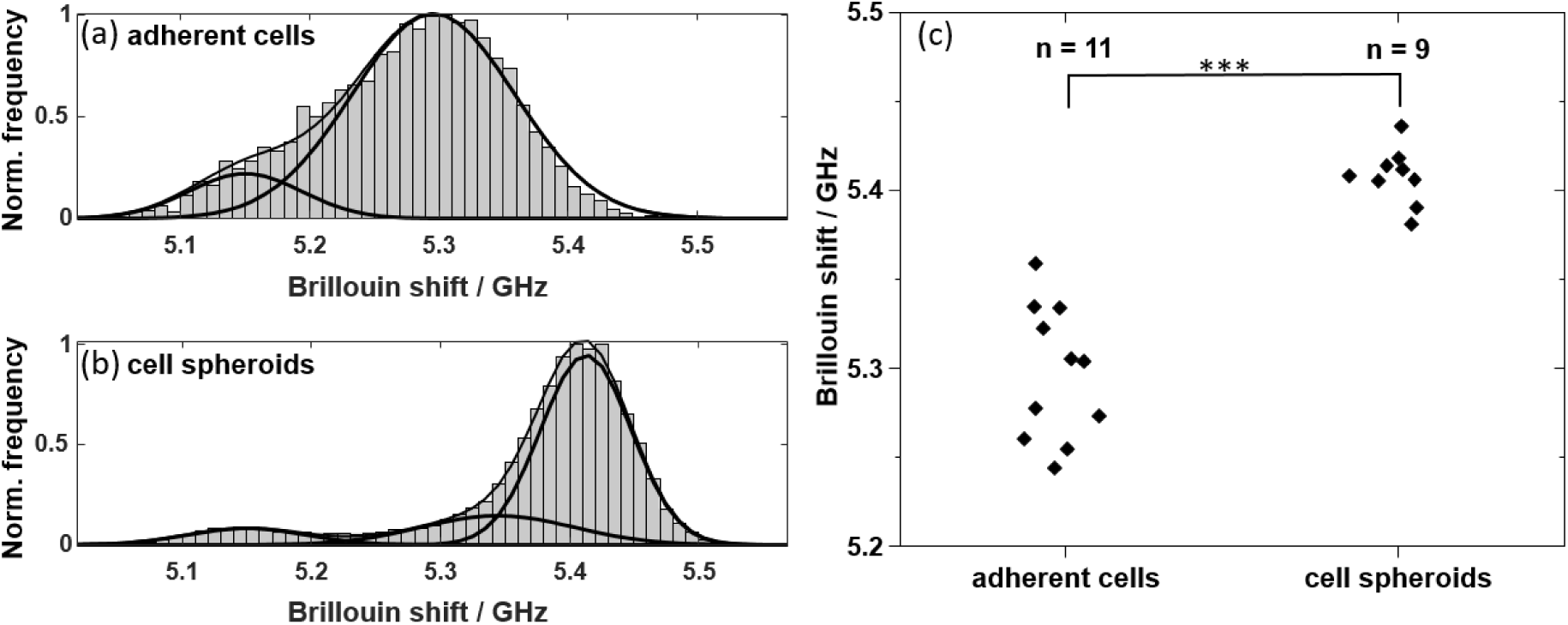
Cumulative frequency histograms (normalized) of the Brillouin shift values of all (a) U87-MG cell (n=11) and all (b) U87-MG spheroid (n=9) mappings showing that there are different contributions fitted by Gaussian curves: Main contributions are at 5.3 GHz and 5.4 GHz, respectively. (c) Brillouin shift frequencies of the main contribution of the individual maps are significantly different (Mann-Whitney U-test; *** p < 0.001).

### 3.3. Combined Brillouin and Raman line scans of U87-MG spheroids highlight biochemical cues underlining the changes of Brillouin shift within spheroids

In order to identify the biochemical reason underlying the higher Brillouin shift that characterizes spheroids in comparison to adherent cells, combined Raman and Brillouin measurements were needed. As the acquisition of Brillouin spectra is about 100 times faster than the acquisition of useful Raman spectra, high resolution maps - as shown in section 3.2 - were only practicable for Brillouin spectroscopy, while for combined measurements the number of measuring points had to be reduced. Therefore, we performed combined Brillouin and Raman line scans across the spheroids, using 15 s integration time, 4 accumulations, 1 μm step size. Two examples of 50 μm long line scans are shown in Figure 5. The Brillouin shift varied in the range of ~5.3 GHz to ~5.55 GHz across the sample (Figure 5a and 5b). A shift in the range ~5.4-5.5 GHz (e.g. in line scan 1 at 9 μm and in line scan 2 at 6 μm) was attributed to the cell body and the Raman spectra were generally characterized by a similar spectral profile as found for cells’ cytoplasm. Regions where the Brillouin shift is close to ~5.3 GHz were interpreted as intercellular spaces (compare with Figure 3b). The highest Brillouin shifts of ~5.5 GHz were observed only in few defined positions. The comparison with Raman spectra at these positions revealed two different spectral patterns (Figure 5c and 5d). One spectral pattern was observed in line scan 1 at 35 μm and in line scan 2 at 32 μm: here the analysis of Raman bands (Figure 5e and 5f) of single spectra allowed clear identification of lipids, based on presence of strong bands at 1267 cm^−1^, 1306 cm^−1^, 1441 cm^−1^, 1659 cm^−1^ and 1748 cm^−1^. In other regions (line scan 1 at 15 μm and line scan 2 at 46 μm), the Raman spectra indicated presence of protein-rich structures based on the bands at 863 cm^−1^, 939 cm^−1^, 1341 cm^−1^, 1462 cm^−1^, 1659 cm^−1^ (50).

**Figure 5.**
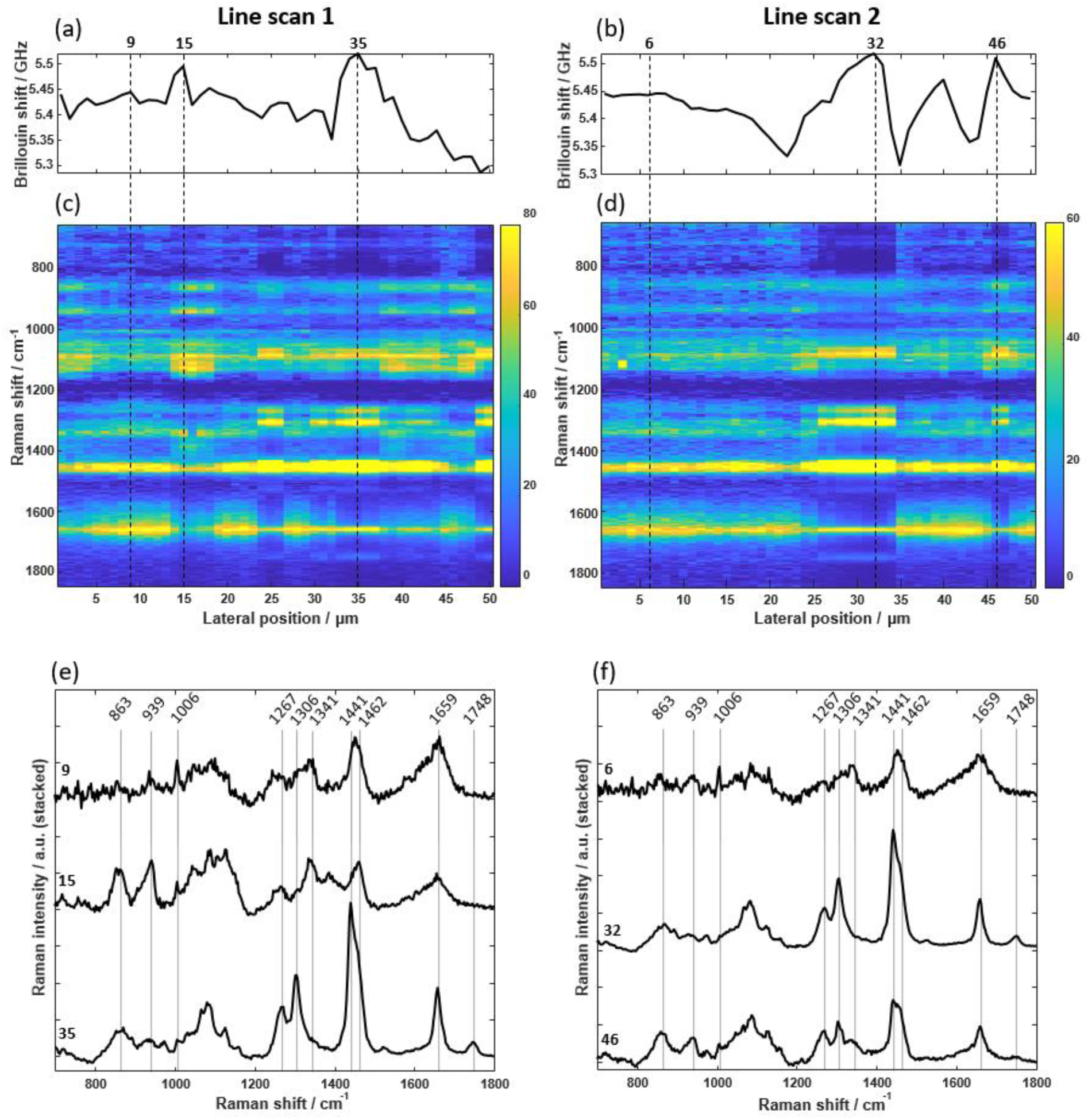
Results of combined line scans on U87-MG spheroids. Brillouin shift profiles (a, b) show regions with high frequencies which can be correlated with spectral patterns visible in the Raman heat maps (c, d), in which the area-normalized Raman intensity is color-coded. Single Raman spectra at specific positions can be attributed to cytoplasm, lipids and protein-rich structures (e, f). Note that the Raman spectrum at 46 μm in line scan 2 actually shows a linear combination of lipid and protein bands indicating the presence of both within the measuring volume.

In order to confirm the presence of lipid accumulation within the spheroids, CARS microscopy was used on fixed spheroids after in vivo spectroscopic measurement. CARS images (n = 7) revealed in fact the presence of intracellular lipid droplets with dimension of few micrometers (Figure 6a), thus compatible with the results of line scans. A quantification (Figure 6b) revealed that lipid droplets accounted for a mean average of 0.58% of the imaged area, which is compatible with the amount of measurement points characterized by a Brillouin shift close or above 5.5 GHz (compare with histogram in Figure 4b).

**Figure 6.**
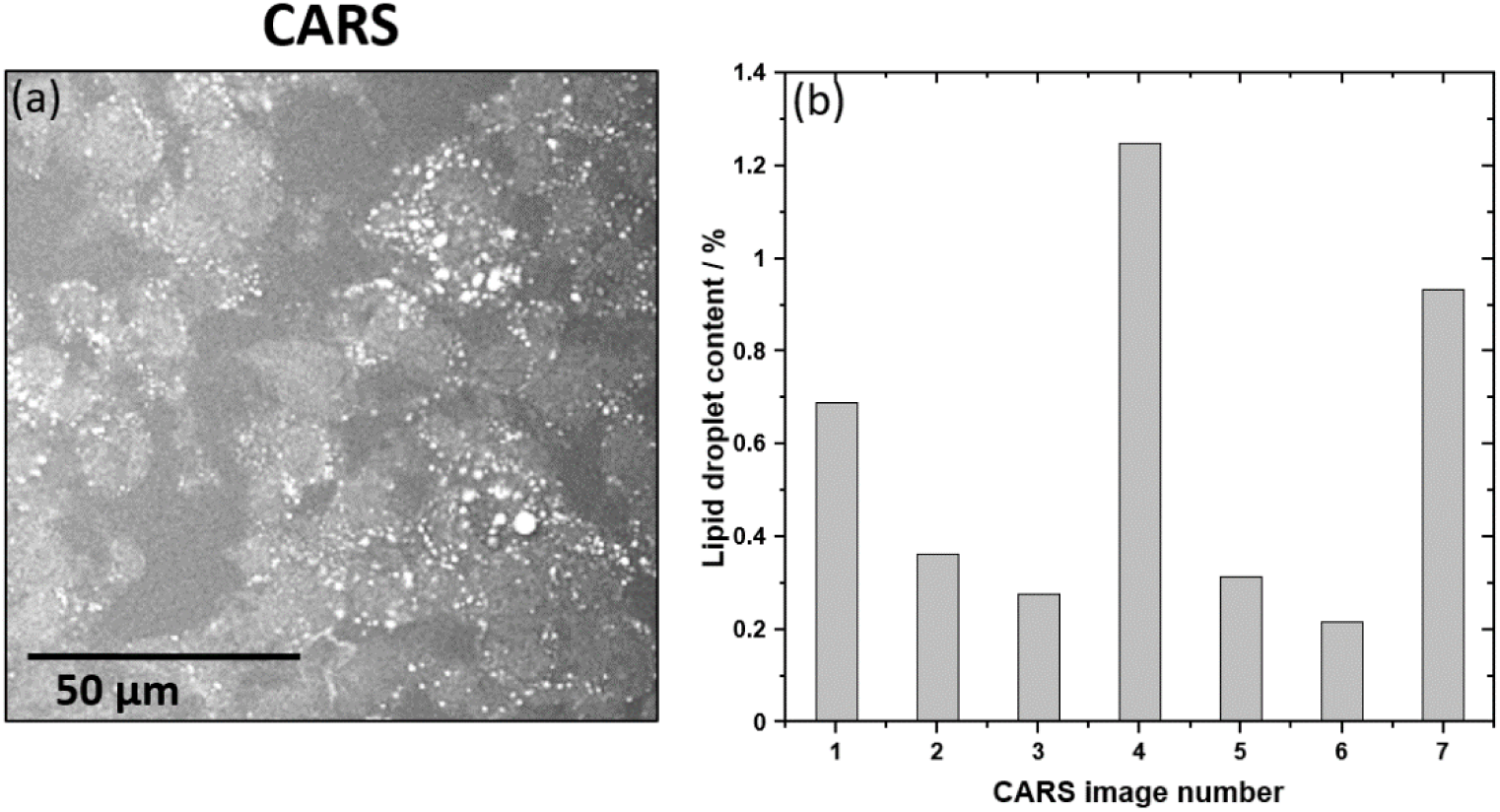
(a) Exemplary CARS image of a U87-MG spheroid cryosection revealing that there are several lipid droplets within the spheroids. (b) Quantification of the lipid droplet content for n = 7 CARS images revealed values between 0.2% and 1.3%, whereby the CARS image in (a) corresponds to image number 1.

## 4. Discussion

Brillouin spectroscopy is a rather new territory in medical research and the interpretation of the Brillouin shift as proxy for the stiffness is still matter for investigations. In contrast, Raman spectroscopy is a successfully established technology on the verge of clinical translation.

As Raman and Brillouin scattering are simultaneously generated in the interaction of laser beams and materials, and because they can be spectrally split by a dichroic mirror, it is possible to combine the two measurements using one excitation laser, one confocal Raman microscope and two spectrometers (32,34,54). We exploited the same principles and realized the system with an infrared laser to reduce elastic scattering and absorption by the tissue and avoid potential photodamage. In order to perform fast acquisition of Brillouin spectra with high extinction and low signal losses, we used a two-stage VIPA setup consisting of two VIPAs with different FSRs resulting in an increased contrast (46). In our system we attained very fast acquisition times for Brillouin spectra, which allowed acquisition of high resolution maps on living cells. In addition, the Brillouin and Raman spectra were measured simultaneously so that biochemical and biomechanical data are registered to allow correlative analysis. However, Raman spectroscopy turned out to be the bottleneck for acquisition time. This limited the acquisition of Raman spectra to relatively small maps of single cells or to line scans across spheroids.

The Raman analysis of U87-MG cells allows to distinguish three different compartments; i.e. the cell nucleus, a perinuclear region and the cytoplasm, independently whether the cells are fixed or living. On the other hand, Brillouin analysis on living cells in culture medium showed an increasing Brillouin shift when going from the cell boundary to the center. By using the Raman spectral information of each pixel, a direct correlation to the Brillouin shift was possible. The three clusters attributed to the cell show higher Brillouin shifts than the surrounding culturing medium, which agrees with previous measurements (47). The cell nucleus has a higher Brillouin shift than the cytoplasm. The nucleolus displays the highest Brillouin shift within adherent cells, and increased Brillouin bandwidth, which indicates higher viscosity. These results are fully consistent with former findings on other types of cells (30,48,55).

Comparing the Brillouin shifts of adherent cells and cell spheroids of the same cell line U87-MG revealed significantly higher values for the latter ones. The Brillouin shift difference of about 0.1 GHz is in line with previous reported values of single cells and spheroids of breast cancer cells (MCF-7) (28). Line scans through spheroids showed local variations of the Brillouin shift. Brillouin shifts of about 5.5 GHz were associated with protein-rich structures and lipid droplets. Regions attributed to cytoplasm based on Raman spectra have a Brillouin shift higher than that of the cytoplasm of adherent cells.

Now, the question is which changes of Brillouin shift indicate also a change in stiffness, as the Brillouin shift *ν*_*B*_ depends not only on the longitudinal modulus, but also on the refractive index and the mass density, as defined by the following equation:

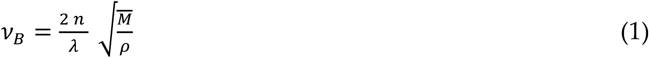

where *n* is the index of refraction, *ρ* the mass density, *M* the longitudinal modulus and *λ* the excitation wavelength. Thus, local variations of *n* and *ρ* may lead to changes in the Brillouin shift even without any change of *M*.

For cytoplasm, nucleus and nucleolus the changes of *n* and *ρ* are expected to compensate each other according to the two-substance mixture model (21,56–58). Therefore, the higher Brillouin shift measured in the cell nucleus and in the nucleolus of cells indicates a higher stiffness of these cellular compartments. Similarly, the higher Brillouin shifts measured in the cytoplasm of spheroid cells compared to adherent cells underline a change of biomechanical properties, i.e. higher stiffness of spheroid cells compared to adherent cells. We attributed the difference to the culturing conditions. This finding demonstrates that the choice of brain tumor model (adherent cells vs. cell spheroids) is highly relevant for analysis of biomechanics and in agreement with other studies on breast cancer spheroids and single cells in hydrogels (28). Future studies on other cell lines may show whether this is a general trend of tumor cells or not.

In case of lipid droplets, the higher Brillouin shift does not indicate stiffness higher than the surrounding cytoplasm: the mass density of lipids is lower compared to cytoplasm (*ρ* = 930 kg/m^3^ (59) vs. *ρ* ≈ 1000 kg/m^3^ (60)) while *n* is higher (*n* = 1.41 (21) vs. n = 1.375 (61)), so that the two contributions do not compensate and the ratio 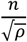 is about 6% higher for lipids compared to the cytoplasm. Indeed, the

Brillouin shift of lipid droplets measured in our experiments (~5.5 GHz) is approximately 5% higher than that of the cytoplasm of adherent cells (5.22 GHz). This result agrees with previous studies on adipocytes, where lipid droplets display a Brillouin shift higher than cytoplasm (21,62), but a 10% lower stiffness by correcting the results for the index of refraction co-registered by optical diffraction tomography (21).

The presence of lipid droplets in glioblastoma cell culture is not surprising, as it is known that they exhibit abnormal lipid metabolism, which plays an important role in aggressiveness (63,64). Lipid droplets function as energy reservoir for glioblastoma and are consumed to support survival by decreased glucose levels (65). This might explain the absence of lipid droplets in the adherent cells, which are known to express lipid droplets mainly during hypoxia while they are missing under normoxic conditions (66,67). Whether there are mechanisms by which cytoplasmic lipid droplets alter glioma cell biomechanics has not been investigated yet. On the other side, lipids possess a clear spectroscopic signature in Raman spectroscopy and their presence and localization can be readily recognized with this technique, thus providing a well-suited reference tool for future investigation of biomechanics of brain tumors.

## 5. Conclusions

Combined Brillouin and Raman spectroscopy proved to be a powerful tool for investigating the interplay between biochemistry and biomechanics of glioblastoma cell cultures. The subcellular resolution of the optical system allowed for a detailed mechanical analysis of adherent cells, whereby the biochemical fingerprint from Raman spectroscopy enabled a correlation of biomechanical properties to cell compartments. The nucleus and especially the nucleolus possess different viscoelastic properties compared to cytoplasm. Moreover, culturing conditions, which result in different cellular architecture and appearance, have an impact on biomechanics, i.e. stiffness is significantly higher in spheroid than in adherent cells, which emphasizes the importance of choosing the appropriate tumor model in future investigations on brain tumor biomechanics. The availability of a co-localized biochemical information obtained by Raman spectroscopy enabled discerning stiffness-related changes of the Brillouin shift as well as localization of stiffness changes at subcellular level. The availability of co-registered biochemical information is very important for a correct interpretation of the biomechanical data on multicellular systems, in particular in the heterogeneous structure of a brain tumor environment. This heterogeneity may also be addressed in future studies e.g. by investigating the microenvironment of spheroids.

## Author Contributions

Conceptualization, M.K., O.U and R.G.; methodology, J.R., O.U. and R.G.; software, J.R.; validation, O.U., R.G.; formal analysis, J.R., O.U. and R.G.; investigation, J.R., K.K. and R.G.; resources, E.K. and G.S.; data curation, J.R. and K.K.; writing—original draft preparation, J.R. and R.G.; writing—review and editing, O.U. and R.G.; visualization, J.R. and R.G..; supervision, E.K., O.U. and R.G..; funding acquisition, M.K. All authors have read and agreed to the published version of the manuscript.

## Institutional Review Board Statement

Not applicable.

## Informed Consent Statement

Not applicable.

## Data Availability Statement

Datasets of this study are available without restrictions on the Open Science Framework: https://osf.io/3xn9h/?view_only=435d50723f784426bb1f4e84b1ee5459

