## Supporting information for "Correlation of biomechanics and cancer cell phenotype by combined Brillouin and Raman spectroscopy of U87-MG glioblastoma cells"

In this document, additional information is provided on "Correlation of biomechanics and cancer cell phenotype by combined Brillouin and Raman spectroscopy of U87-MG glioblastoma cells".

### 1. Exemplary Brillouin spectrum with Lorentzian fitting

In order to demonstrate the non-linear fitting procedure, we present here an exemplary Brillouin spectrum with fitted Lorentzian functions (blue curves). The two bands at ±5.3 GHz correspond to the sample signal, the two bands at ±3.81 GHz belong to the methanol reference and the two bands in the center are the shoulders of the strongly filtered Rayleigh signal. The sum of all six Lorentzian functions (red curve) fits the measuring points (black dots).

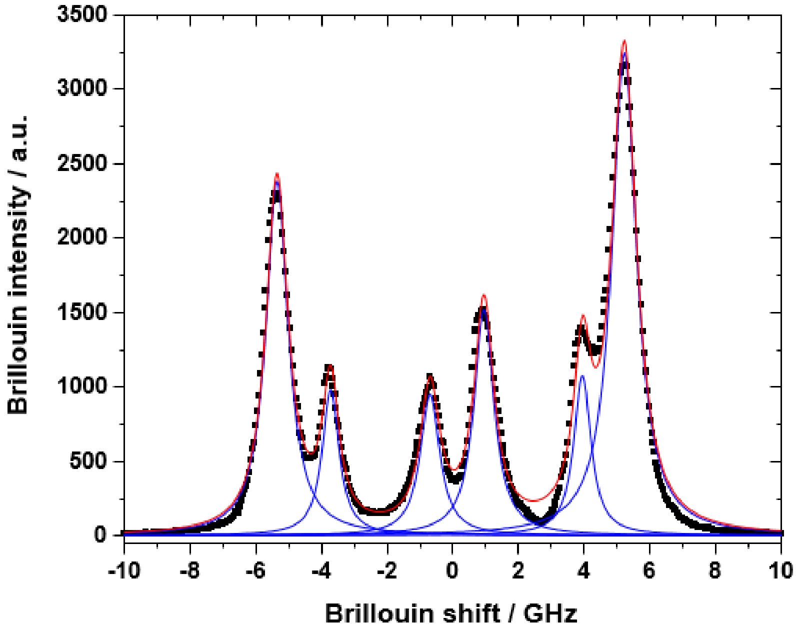


**Figure S1.** Exemplary Brillouin spectrum fitted with Lorentzian functions (blue curves), so that their sum (red curve) best fits to the measuring points (black dots).

**2. Removal of the culturing medium contribution in the cell histograms**

Because the cells' Brillouin maps consist to a large extent of surrounding culturing medium and not only of the cells itself, the corresponding histograms encompass a large medium contribution (at 5.1-5.2 GHz), which is not of interest for further analysis. Therefore, we decided to fit this contribution with a Gaussian function (Figure S2a) and subtract it from the histogram. The remaining histogram (Figure S2b) then shows only the contribution of interest, i.e. the cell's contribution. Another fitting determines its mean Brillouin shift. It should be mentioned that the subtraction of the medium contribution does not lead to different results regarding the cell contribution, but is performed only because of visualization reasons.

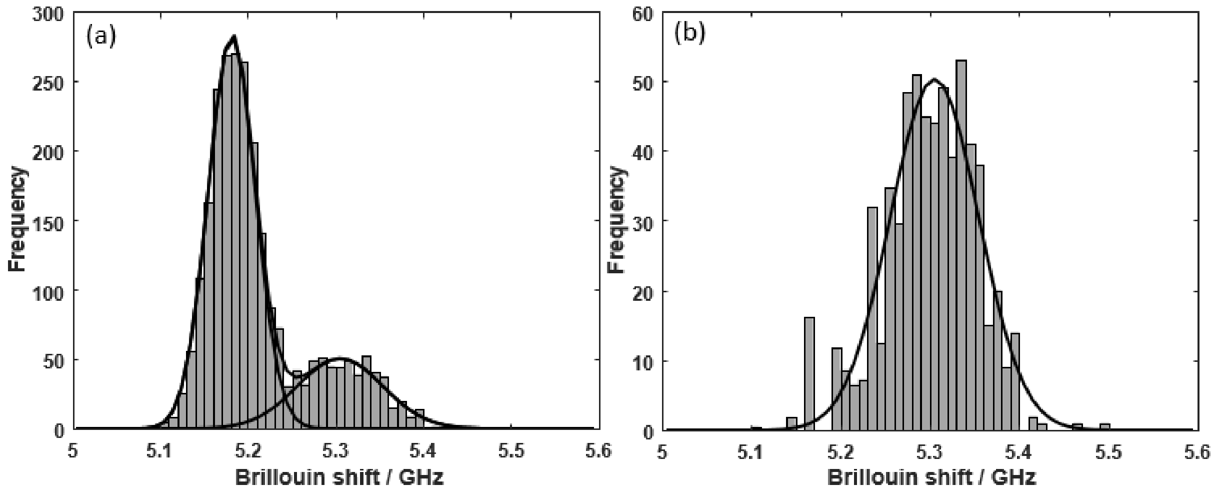


**Figure S2.** (a) Exemplary Brillouin shift histogram of an adherent cell measurement showing a major culturing medium contribution at 5.1-5.2 GHz. (b) Same histogram as in (a), but with the medium contribution subtract, indicating that the mean cell contribution is around 5.3 GHz.

**3. Additional combined measurements of living U87-MG cells**

In addition to the U87-MG cell measurement presented in Figure 2 of the main manuscript, we would like to show three more examples, which were acquired in same manner. They indicate the robustness of the cluster analysis, which was always able to identify the same subcellular structures. Therefore, the centroid Raman spectra are similar to those presented in the main manuscript. Moreover, the Brillouin maps and the Brillouin shift dot plot show same characteristics as discussed in section 3.1., i.e. the Brillouin shift of the nucleus/nucleolus is highest, the nucleolus shows the broadest Brillouin linewidth and the Brillouin intensity is behaving in an inverse manner. Note that sometimes the upper part of the Brillouin intensity map has significantly higher values (e.g. see first 2-3 lines in Figure S3.1d), which is due to focus issues at the beginning of each measurement, but not due to the cell.

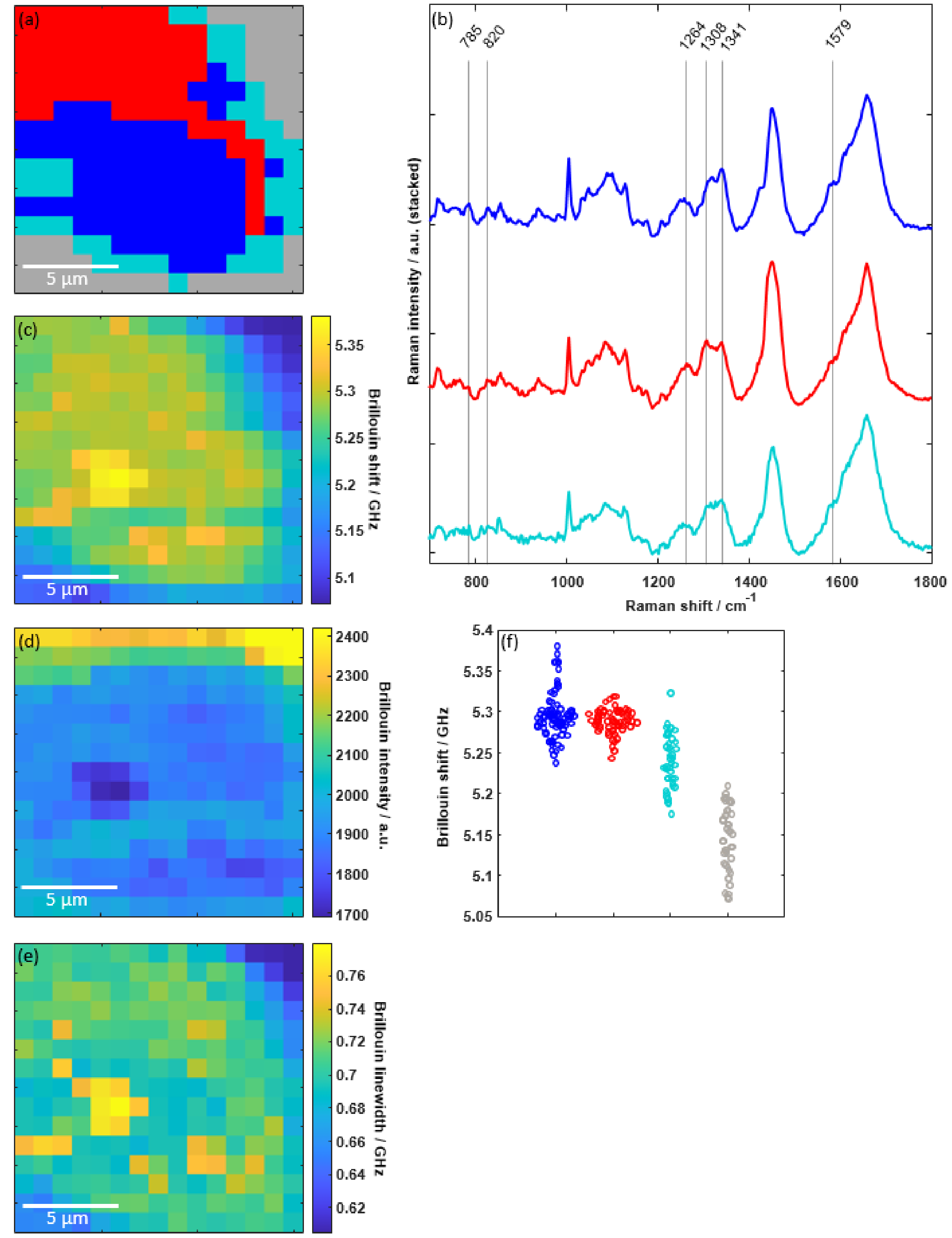


**Figure S3.1** (a) Raman cluster map of a U87-MG cell consisting of four different clusters. (b) Mean Raman spectra of the three clusters associated with the cell being the nucleus (blue), perinuclear region (red) and the cytoplasm (cyan). Simultaneously acquired Brillouin shift (c), Brillouin intensity (d) and Brillouin linewidth (e) map revealing all three the cellular structure. (f) Brillouin shift values for each pixel are assigned to the respective cell compartment obtained by the cluster analysis of Raman spectra.

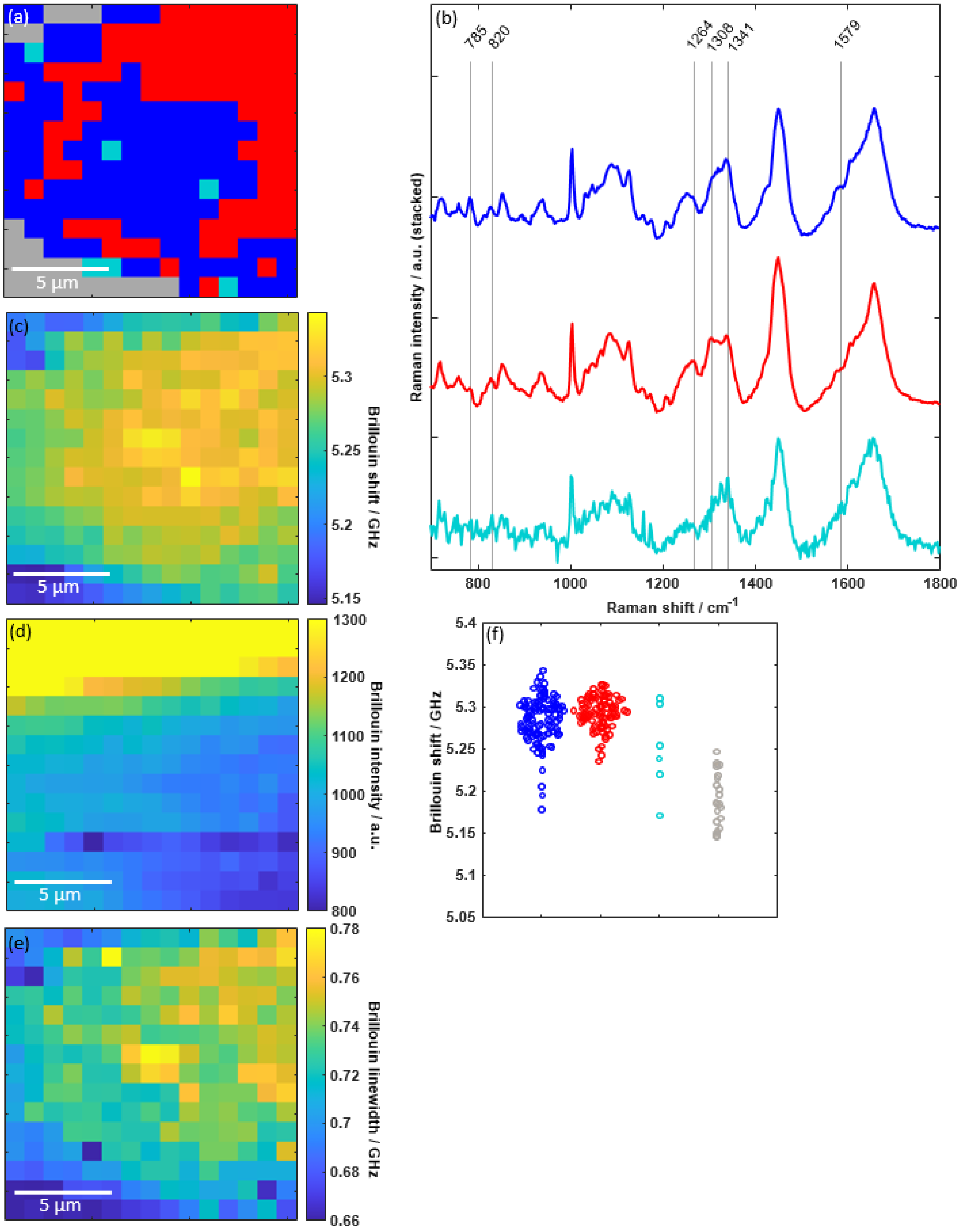


**Figure S3.2** (a) Raman cluster map of a U87-MG cell consisting of four different clusters. (b) Mean Raman spectra of the three clusters associated with the cell being the nucleus (blue), perinuclear region (red) and the cytoplasm (cyan). Simultaneously acquired Brillouin shift (c), Brillouin intensity (d) and Brillouin linewidth (e) map revealing all three the cellular structure. (f) Brillouin shift values for each pixel are assigned to the respective cell compartment obtained by the cluster analysis of Raman spectra.

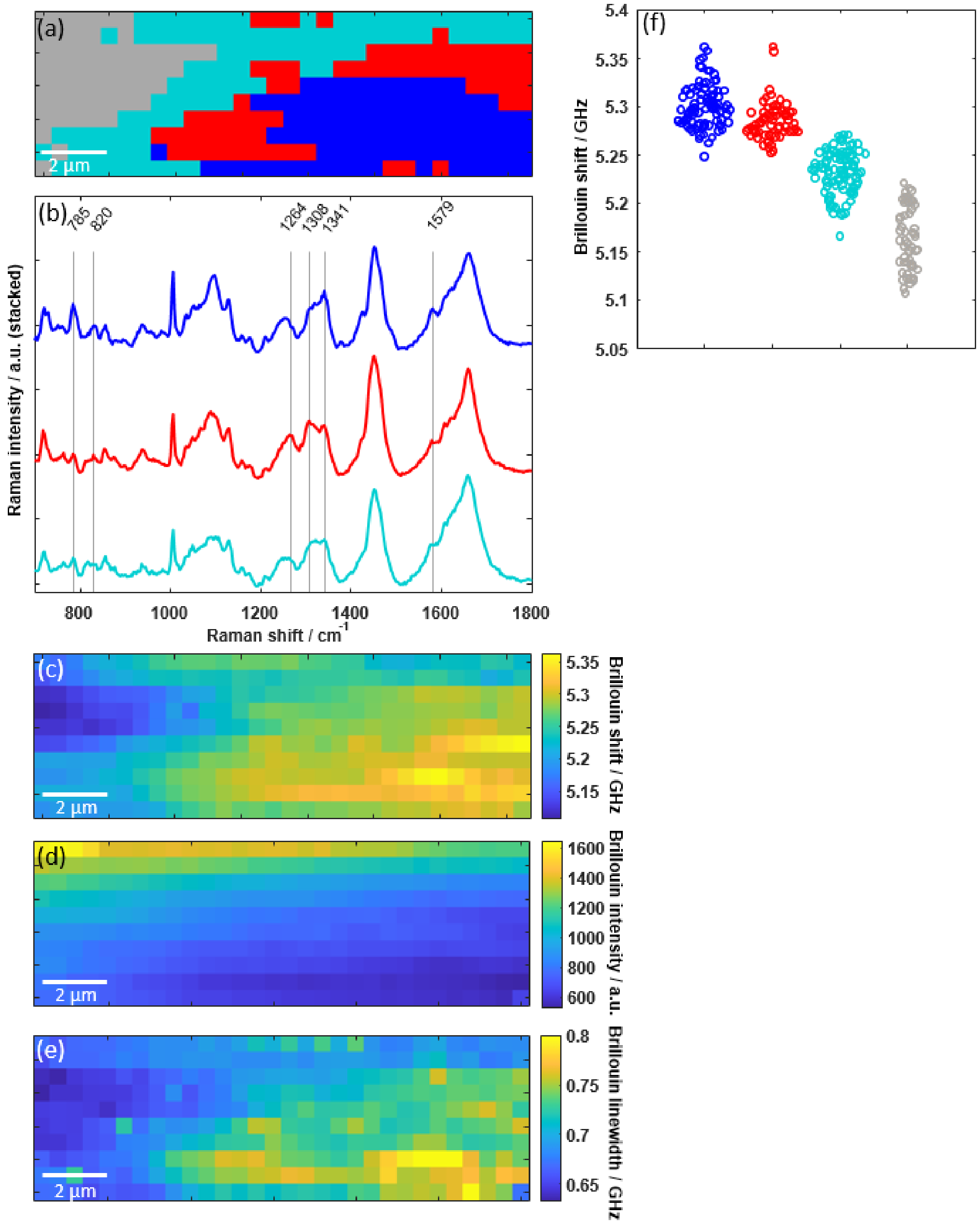


**Figure S3.3** (a) Raman cluster map of a U87-MG cell consisting of four different clusters. (b) Mean Raman spectra of the three clusters associated with the cell being the nucleus (blue), perinuclear region (red) and the cytoplasm (cyan). Simultaneously acquired Brillouin shift (c), Brillouin intensity (d) and Brillouin linewidth (e) map revealing all three the cellular structure. (f) Brillouin shift values for each pixel are assigned to the respective cell compartment obtained by the cluster analysis of Raman spectra.

**4. Fixed U87-MG cell**

In order to verify the quality of the Raman spectra clustering and the assignment of the mean spectra to the cell compartments, Raman maps of n = 8 formalin fixed U87-MG cells were acquired. The fixation allows for longer measuring times compared to maps of living cells, wherefore a higher amount of pixel per map can be chosen, which increases the resolution of the map. Additionally, the acquisition time of each single spectrum can be increased resulting in an increased signal-to-noise ratio. Figure S4a shows an exemplary Raman map of a fixed cell (25 s integration time, 4 accumulations, 1 μm step size) consisting of four clusters, again. Note that here the gray cluster is not associated with surrounding culturing medium but with surrounding air. The clustering map reveals the same structures visible in the bright field image (Figure S4b). The mean spectra of the four clusters are depicted in Figure S4c. The Raman spectra of the three clusters associated with the cell are in great accordance with those retrieved from living cells (Figure 2b of the main manuscript) and thus confirm the attribution to the cell compartments.

To further validate the cluster assignment, we performed hematoxylin and eosin (HE) staining of the same fixed U87-MG cells. The HE staining (Figure S4d) clearly resolves the nucleus. Its location matches very well with the blue cluster, wherefore its assignment to the nucleus is straightforward. A clear difference of the perinuclear region (red cluster) and the cell periphery, i.e. the remaining cytoplasm (cyan cluster), cannot be appreciated by the HE staining. However, these two regions are different in lipid content, as the endoplasmic reticulum localize in a lipid-rich perinuclear region, in contrast to the cell periphery, which has less lipids and proportionally more proteins. This difference was visualized by CARS microscopy on a similar cell grown under same conditions. The CARS intensity image (Figure S4e) confirms inhomogeneous lipid distribution within the cell and thus encourages the assignment of the red cluster to a lipid-rich perinuclear region and the cyan cluster to the remaining cytoplasm.

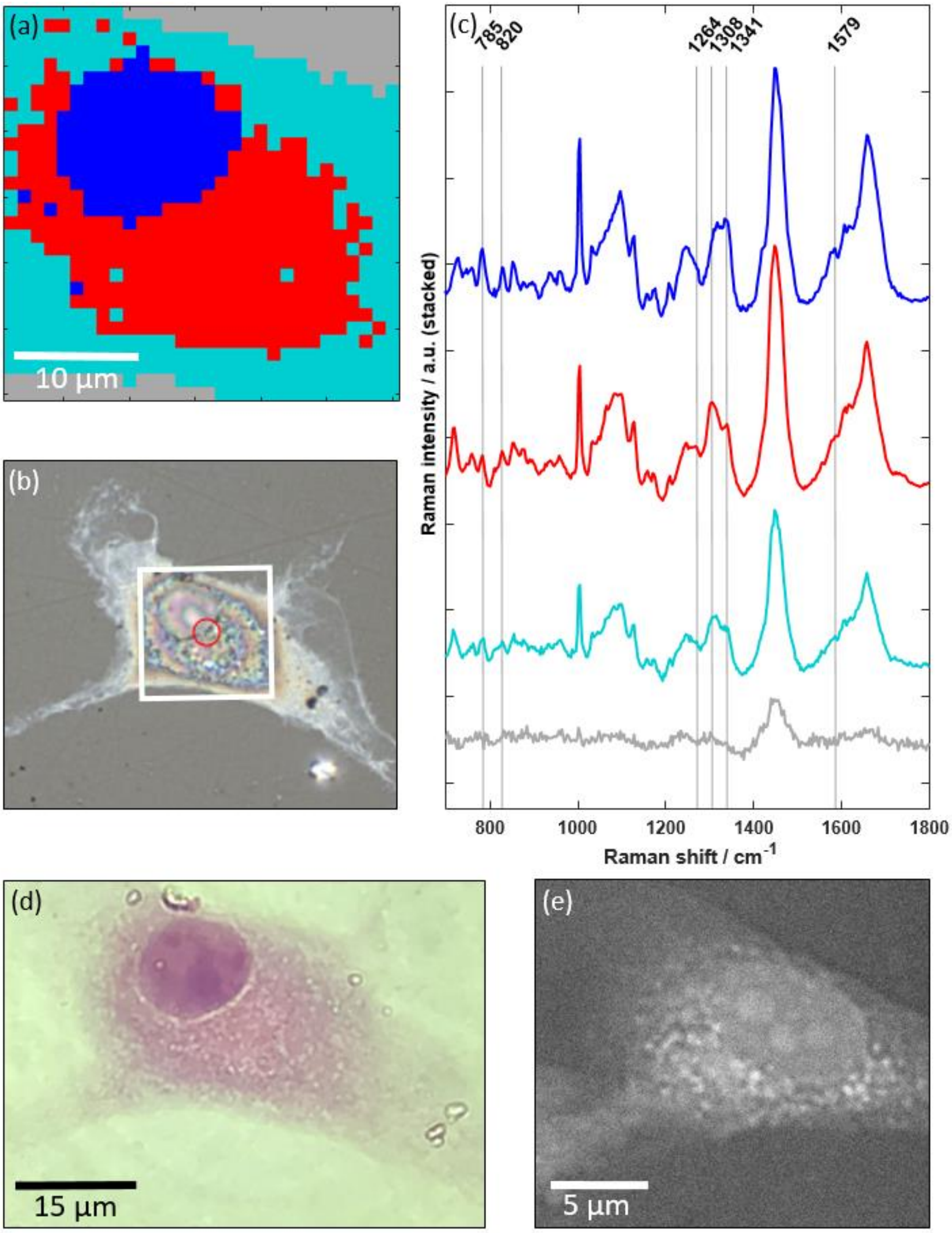


**Figure S4.** (a) Raman cluster map of a formalin fixed U87-MG cell (bright field image in (b); white box is 30 × 30 µm²) and (c) the corresponding mean spectra being in great accordance to that shown in Figure 2b of the main manuscript. (d) HE staining of the same cell clearly resolving the nucleus, being associated with the blue cluster. The different lipid content of the perinuclear (red) cluster and the remaining cytoplasm (cyan) cluster can be seen in the CARS intensity image (e) of a similar cell grown under same conditions.
